# Pathogenic and transcriptomic differences of emerging SARS-CoV-2 variants in the Syrian golden hamster model

**DOI:** 10.1101/2021.07.11.451964

**Authors:** Kyle L. O’Donnell, Amanda N. Pinski, Chad S. Clancy, Tylisha Gourdine, Kyle Shifflett, Paige Fletcher, Ilhem Messaoudi, Andrea Marzi

## Abstract

Following the discovery of severe acute respiratory syndrome coronavirus 2 (SARS-CoV-2) and its rapid spread throughout the world, new viral variants of concern (VOC) have emerged. There is a critical need to understand the impact of the emerging variants on host response and disease dynamics to facilitate the development of vaccines and therapeutics. Syrian golden hamsters are the leading small animal model that recapitulates key aspects of severe coronavirus disease 2019 (COVID-19). In this study, we show that intranasal inoculation of SARS-CoV-2 into hamsters with the ancestral virus (nCoV-WA1-2020) or VOC first identified in the United Kingdom (B.1.1.7) and South Africa (B.1.351) led to similar gross and histopathologic pulmonary lesions. Although differences in viral genomic copy numbers were noted in the lungs and oral swabs of challenged animals, infectious titers in the lungs were comparable. Antibody neutralization capacities varied, dependent on the original challenge virus and cross-variant protective capacity. Transcriptional profiling indicated significant induction of antiviral pathways in response to all three challenges with a more robust inflammatory signature in response to B.1.1.7. Furthermore, no additional mutations in the spike protein were detected at peak disease. In conclusion, the emerging VOC showed distinct humoral responses and transcriptional profiles in the hamster model compared to the ancestral virus.

## Introduction

Severe acute respiratory syndrome coronavirus-2 (SARS-CoV-2) has emerged as a novel, highly infectious respiratory CoV and the causative agent of CoV disease 2019 (COVID-19)^1^. First described in the city of Wuhan in Hubei province of China, SARS-CoV-2 is a member of the *Coronavirdae* family, which possess large, non-segmented RNA genomes^1^. High levels of transmission, especially in regions with low vaccination rates, facilitate the emergence of mutations that improve viral fitness. SARS-CoV-2 variants of concern (VOC) are defined as variants that have one or more mutations that confer worrisome epidemiologic, immunologic, or pathogenic properties^2^. Several SARS-CoV-2 VOC have emerged such as B.1.1.7 first reported in the United Kingdom (UK), which is associated with increased transmission compared to the ancestral virus reported from Washington, USA in early 2020^3^. This variant acquired over 20 mutations including N501Y within the spike (S) protein that increased binding affinity to the angiotensin converting enzyme 2 (ACE2) receptor^4,5^. In addition, the S protein of the B.1.1.7 variant has a deletion of amino acids 69 and 70 which has been shown to increase viral escape in immunocompromised individuals^6,7^. VOC B.1.351 was originally reported in South Africa (SA) and harbors similar mutations in S compared to B.1.1.7 as well as the K417N and E484K substitutions that may decrease the efficacy of existing vaccines^8–12^. Other variants more recently reported in the United States (B.1.427, B1.429) also harbor mutations in S (e.g., N501Y) that have been associated with reductions in neutralizing antibody titers^13^.

There is an urgent need to understand the effect of new mutations within VOC on the host immune response to facilitate the development of vaccines and therapeutics. In this study, we compared pathologic features of and immune responses to the original virus (ancestral), and the later B.1.1.7 and B.1.351 variants in the well-established Syrian golden hamster model of severe COVID-19^14^. Specifically, we longitudinally assessed viral replication, histopathological changes, development of humoral immunity and humoral cross-reactivity amongst VOC. Additionally, we employed RNA-seq and digital cell quantification of lung homogenates to determine differences in transcriptomic signatures and to infer changes in immune cell subsets. We identified similar histopathological changes, levels of infectious virus, and antibody titers amongs all infections. However, transcriptional responses and the capacity to cross-neutralize SARS-CoV-2 was VOC-dependent. Collectively, these data demonstrate that mutations within SARS-CoV-2 modulate host defense pathways.

## Results

### Gross lung pathology

Syrian golden hamsters were separated into three cohorts (n=15 per cohort) and challenged intranasally (IN) with 10^5^ TCID_50_ of one of three different SARS-CoV-2 variants: ancestral (nCoV-WA1-2020), B.1.1.7, and B.135. Five uninfected animals served as negative controls. Scheduled necropsies were performed at 4, 14, and 28 days post-challenge (DPC) for all groups to capture peak disease and convalescence (Fig. S1A). Peak weight loss was achieved amongst all three groups 7 DPC, however, no significant difference in body weight changes occurred over the first 10 DPC for any of the infections (Fig. S1B). Gross pulmonary lesions were observed in all infected hamsters at 4 DPC (Fig. S1D). Lungs harvested 4 DPC showed multifocal to locally extensive areas of red to purple coloration (consistent with consolidation) disseminated throughout all lung lobes. Additionally, lungs generally failed to collapse indicating interstitial disease. Lung samples harvested 14 and 28 DPC had either no gross lesions or limited, small, multifocal areas of consolidation and/or congestion. Analysis of histopathology samples demonstrated evidence of interstitial pneumonia on 4 and 14 DPC in all groups (Fig. S1C).

### Histopathology and immunohistochemistry of hamster lungs

Pulmonary pathology consistent with previously described coronavirus respiratory disease was observed at 4 DPC in lung samples from hamsters infected with each virus (Fig. 1)^15^. Five uninfected animals served as negative controls (Fig. 1A, E, I). Foci of interstitial pneumonia and bronchiolitis were observed throughout all evaluated lung lobes of infected hamsters. Minimal to mild bronchiolitis characterized by individual epithelial cell necrosis, epithelial cell basophilia and hyperplasia and rare syncytial cell formation was observed throughout all variants (Fig. 1 B-D). Interstitial pneumonia varying in percent of lung involvement and moderate to severe severity was observed within each animal regardless of the variant. Interstitial pneumonia at 4 DPC was defined by expansion of alveolar septa by edema fluid, leukocyte infiltration and fibrin, with leukocyte spillover into adjacent alveolar spaces and in severe cases, complete loss of pulmonary architecture (Fig. 1 F-H). Tracheitis characterized by neutrophilic influx and epithelial cell necrosis was observed in all evaluated sections of trachea in each animal at 4 DPC. Immunohistochemical analysis showed immunoreactivity to an antibody specific to SARS-CoV-2 within bronchiolar epithelia, type I and type II pneumocytes and macrophages in lungs of all hamsters regardless of the viral variant (Fig. 1 J-L).

**Fig. 1.**
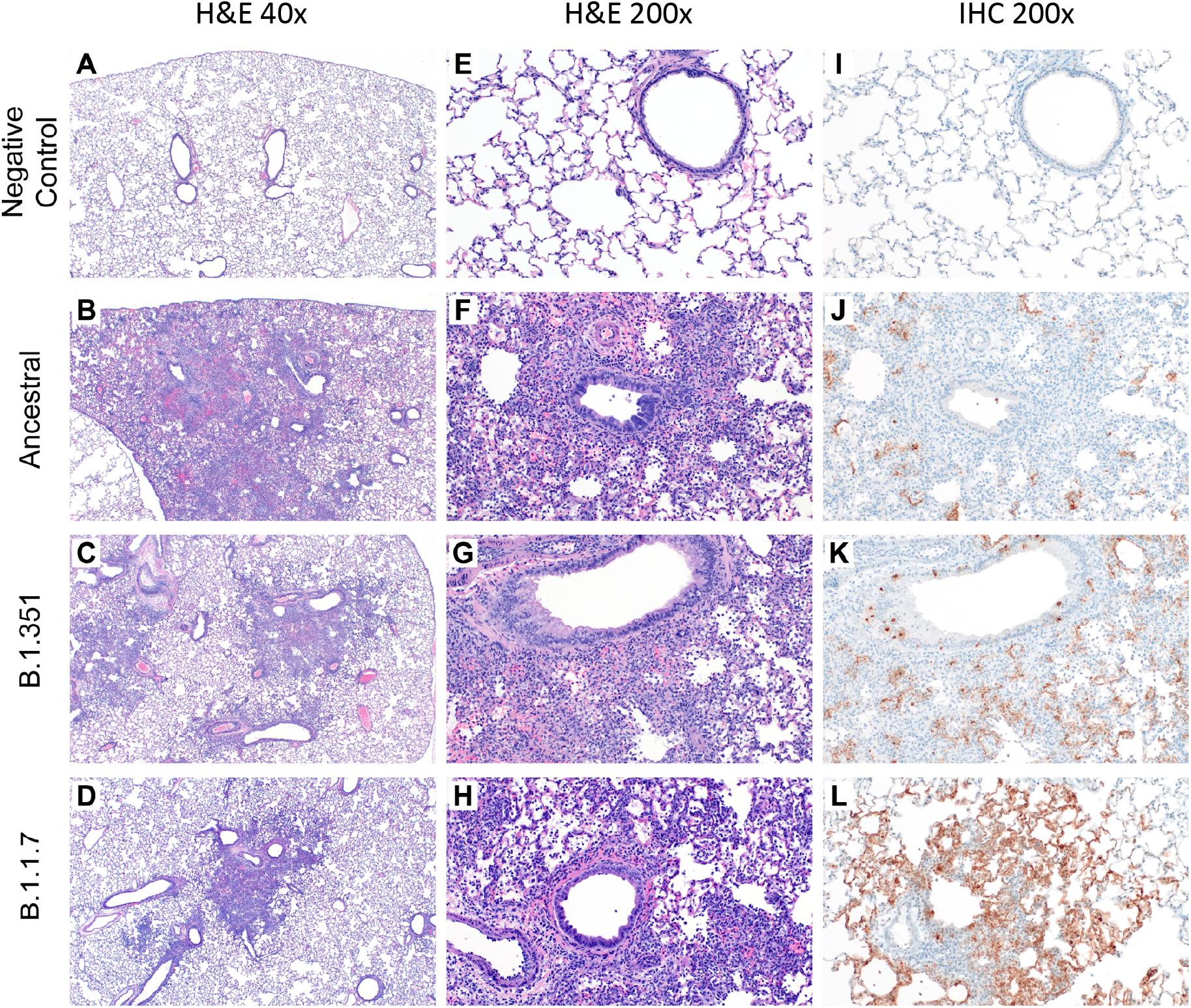
Histopathology and Immunohistochemistry of hamster lungs. **(A-H)** Representative H&E images of lungs of hamsters infected with 10^5^ TCID_50_ of ancestral, B.1.1.7, and B.1.351 variants at 4 days post-challenge (DPC). Foci of interstitial pneumonia and bronchiolitis were observed throughout all evaluated lung lobes of infected hamsters. **(I-L)** Immunohistochemistry (IHC) detected SARS-CoV-2 nucleocapsid staining in the lungs of all infected hamsters.

At 14 and 28 DPC pulmonary pathology was similar in lungs of hamsters infected with all viruses (Fig. S1C, D). Foci of persistent type II pneumocyte hyperplasia with occasional apical cilia formation (alveolar bronchiolization) adjacent to terminal bronchioles was observed throughout all lung lobes. Frequently, foci of alveolar bronchiolization entrapped low to moderate numbers of foamy macrophages. Antigen was not detected by immunohistochemical evaluation for any viral variant at either 14 or 28 DPC.

### Viral burden

Total viral RNA copy numbers and infectious viral titers were quantified in lungs of challenged animals at the three time points mentioned above (Fig. 2A-C). There was no difference in viral RNA copy numbers amongst challenged groups at 4 DPC (Fig. 2A). However, there was significantly more viral RNA at 14 DPC in the B.1.1.7-challenged group compared to the ancestral and B.1.351 groups. At 28 DPC there were significantly more viral RNA copies in the lungs of ancestral-challenged hamsters than the B.1.1.7 group (Fig. 2A). We also assessed sub-genomic viral RNA (sgRNA) as a surrogate of active viral replication^16,17^. Levels of lung sgRNA peaked at 4 DPC and were comparable among the three variants (Fig. 2B). In contrast, we observed a significant difference in sgRNA among the groups at 14 DPC. Specifically, B.1.1.7-infected hamsters exhibited the highest residual sgRNA present compared to the ancestral and B.1.351 groups (Fig. 2B). The B.1.351 group also had significantly higher sgRNA levels compared to the ancestral group (Fig. 2B). However, infectious viral titers were only detected 4 DPC in lungs in all hamsters (Fig. 2C).

**Fig. 2.**
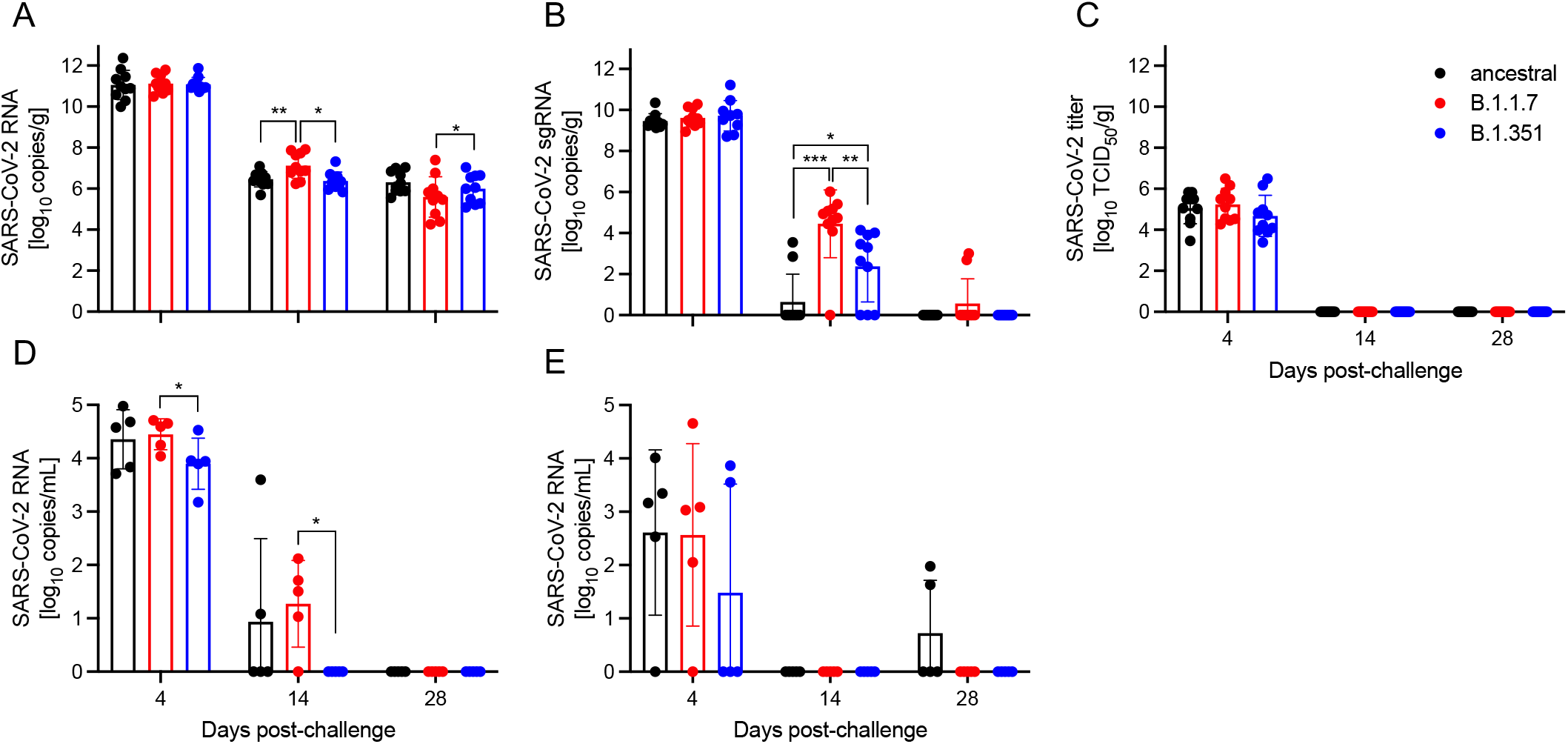
SARS-CoV-2 burden in lungs, oral swabs and blood. **(A)** Total SARS-CoV-2-specific RNA and **(B)** total SARS-CoV-2-specific sub-genomic RNA (sgRNA) in the lungs of challenged animals at 4, 14, and 28 days post-challenge (DPC). **(C)** Infectious SARS-CoV-2 titer in the lungs of infected hamsters. Total SARS-CoV-2-specific RNA in the **(D)** oral swabs and **(E)** blood of infected hamsters at the time of euthanasia. Geometric mean and standard deviation (SD) are depicted; statistical significance is indicated ***p < 0.001, **p <0.01 and *p < 0.05.

Oral viral shedding and viremia were evaluated at the time of necropsy. Infection with the B.1.1.7 VOC resulted in significantly more oral viral shedding than the B.1.351 variant at 4 and 14 DPC (Fig. 2D). Viremia peaked at 4 DPC and was comparable amongst all infections (Fig. 2E). Profiling the viral genomes recovered from the lungs of infected hamsters at 4 DPC revealed no changes in the viral sequences in ancestral and B.1.1.7-infected animals compared to the reference genomes (Table 1). However, we identified three mutations in all B.1.351-infected animals, including two nonsynonymous mutations in 5’ UTR (T201C) and nsp3 (G172C), and one synonymous mutation in nsp3 (G5942G). A single B.1.351-infected animal presented with an additional mutation (L3892F) in nsp3 (Table 1). No mutations in S were detected.

**Table 1.** Genome comparison of SARS-CoV-2 variants.

| Variant of concern | Amino acid changes (# of animals affected, 4 DPC) |
| --- | --- |
| ancestral | none detected |
| B.1.1.7 | none detected |
| B.1.351 | 5' UTR, T201C (5/5)<br>nsp3, L3892F (1/5)<br>nsp3, G5942G (5/5)<br>ORF3a, G172C (5/5) |

### Humoral immune responses post-challenge

We utilized standard ELISA methods to determine the SARS-CoV-2 S-specific IgG responses, and S receptor-binding domain (RBD)-specific IgG responses. There was no difference in the S-specific IgG titers at either 14 or 28 DPC amongst the groups (Fig. 3A). Similarly, no difference was determined in the RBD-specific IgG titers at 14 DPC (Fig. 3B). However, at 28 DPC the RBD-specific IgG titer was significantly higher in animals challenged with B.1.351 compared to B.1.1.7 (Fig. 3B).

**Fig. 3.**
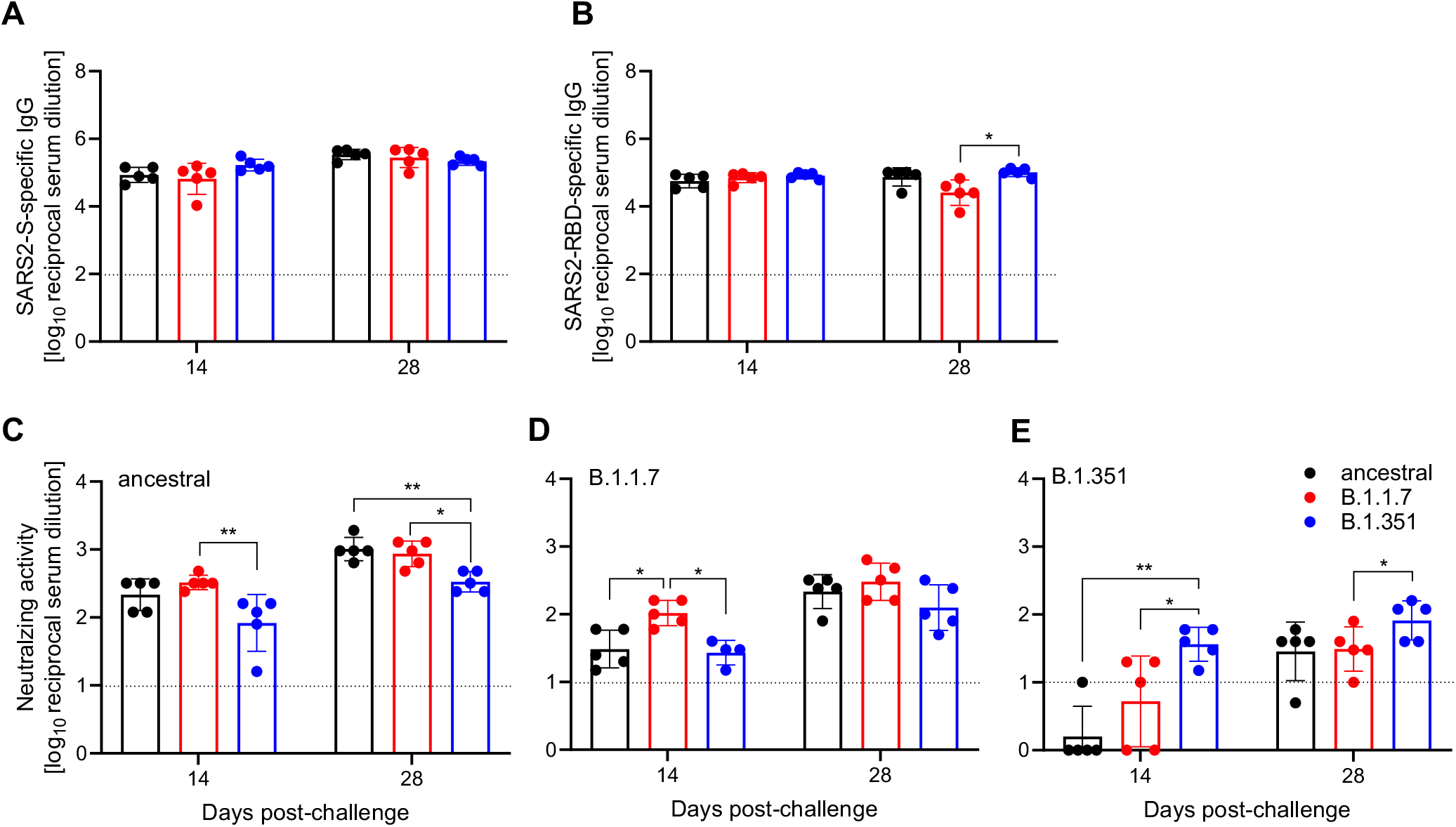
Humoral immune response in challenged hamsters. Serum samples were collected at 14 and 28 DPC and by ELISA. **(A)** Total SARS-CoV-2 S-specific IgG and **(B)** RBD-specific IgG are shown. Cross-variant and homologous neutralization was assessed against **(C)** ancestral, **(D)** B.1.1.7, and (E) B.1.351 viruses. Line indicates limit of detection. Geometric mean and SD are depicted; statistical significance is indicated **p <0.01 and *p < 0.05.

Next, we assessed the functionality of the humoral response by neutralization assay, not only against the homologous challenge virus, but also against the other two variants to determine cross-reactivity generated from the primary infection. Hamsters challenged with the ancestral virus exhibited comparable neutralizing titers against the homologous challenge variant (ancestral) and the B.1.1.7 variant at 14 and 28 DPC but lower titers against the B.1.351 variant at both timepoints assessed (Fig. 3C). In contrast, hamsters challenged with the B.1.1.7 or the B.1.351 variant each exhibited significantly higher neutralizing titers against their homologous challenge virus at 14 DPC compared to variants to which they were not exposed (Fig.3D, E). This difference persisted for the B.1.351-infected animals at 28 DPC when comparing anti-B.135.1 and anti-B.1.1.7 titers (Fig. 3E). Moreover, the overall neutralization titers against the B.1.351 variant were 1-2 logs lower than the other two variants regaradless of homolgous or heterologous assessment.

### COVs elicit unique transcriptional responses in the lungs

To elucidate differences in the host responses to VOC, we profiled the transcriptional responses in lung tissues obtained at peak viral loads (4 DPC) (Fig. 4, S2). Principal component analysis (PCA) revealed distinct separation between uninfected and uninfected animals (Fig. S2A), with the B.1.1.7 variant infection resulting in the most distinct transcriptional profile and the largest number of differentially expressed genes (DEGs) (n=1,277) while infection with B.1.351 resulted in the smallest number of DEGs (n=395) (Fig. 4A-C). Most DEGs were upregulated following infection with all three viruses (Fig. 4A-C). A core of 291 DEGs was shared by all variants and an additional ^~^270 DEGs were shared only between B.1.1.7- and ancestral-infected hamsters (Fig. 4D).

**Fig. 4.**
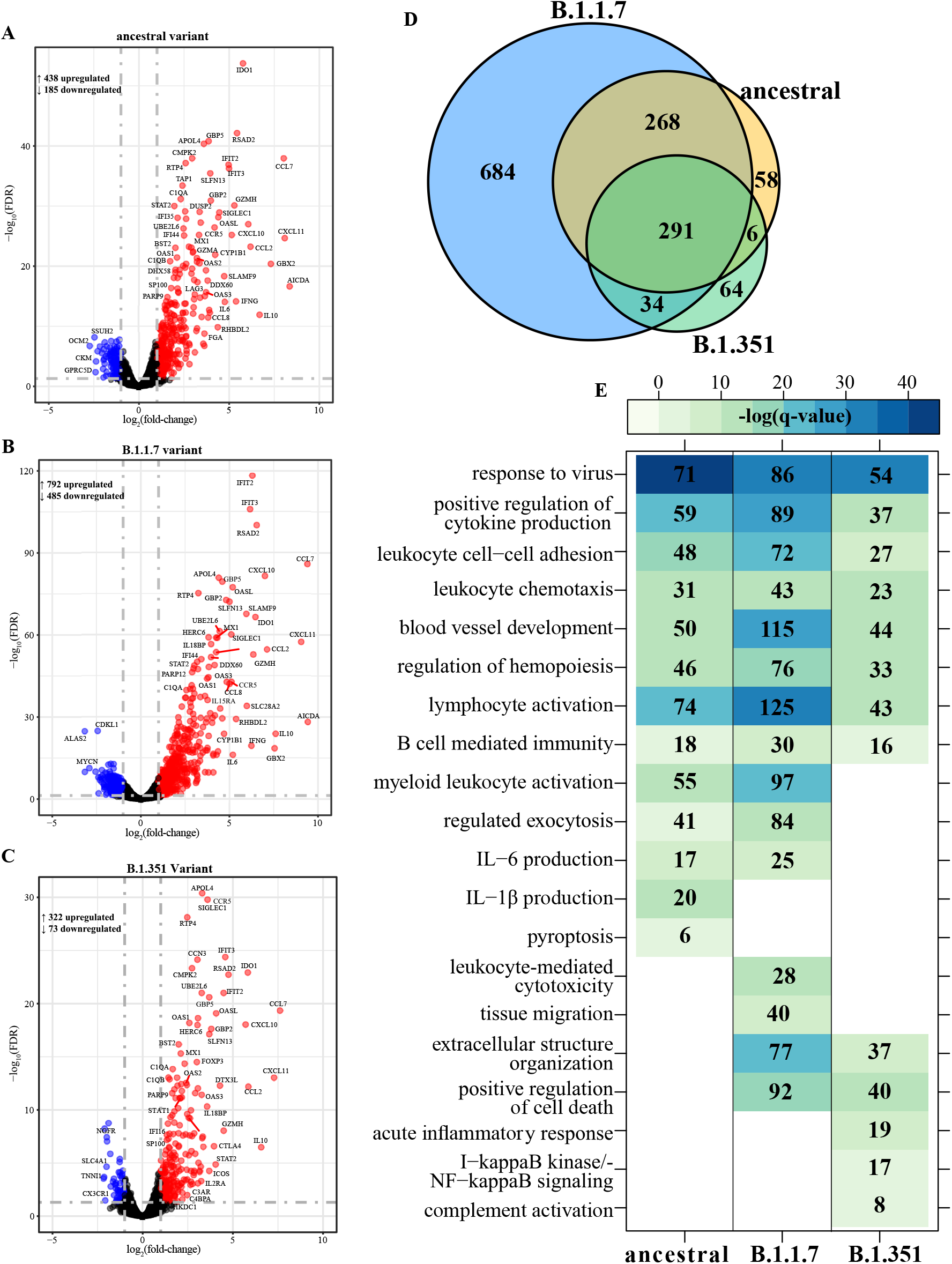
SARS-CoV-2 variants induce distinct transcriptional changes. Volcano plot of global gene expression changes at 4 DPC with SARS-CoV-2 **(A)** ancestral, **(B)** B.1.1.7 or **(C)** B.1.351 variants. Downregulated and upregulated differentially expressed genes (DEGs; average RPKM ≥ 1) are colored blue and red, respectively. Exemplary genes are labeled. **(D)** Venn diagram of DEGs determined in panels A-C. **(E)** Functional enrichment of DEGs determined following each infection in panels A-C. Color intensity represents statistical significance as the negative log of the FDR-adjusted p-value [-log(q-value)], with range of colors based on the GO terms with the lowest and highest statistical value for all GO terms present. Blank boxes indicate no statistical significance. Numbers of DEGs enriching to each GO term are noted in each box.

We performed functional enrichment of DEGs in order to determine their biological relevance. DEGs induced by all three viral infections enriched to Gene Ontology (GO) terms associated with antiviral immunity (e.g., “response to virus”), immune cell recruitment (e.g., “leukocyte chemotaxis”) and mobilization of adaptive immunity (e.g., “lymphocyte activation”, “B cell-mediated immunity”) (Fig. 4E). DEGs enriching to “response to virus” and common to all three infections play roles in type I interferon (IFN) signaling (e.g., *IRF7, IRF9, STAT1/2*), nucleic acid detection (e.g., *DDX60, DHX58*) and the antiviral response (e.g., *ISG15, MX1, RSAD2, SAMHD1*) (Fig. S2B). These DEGs were upregulated following infection with all three variants, particularly B.1.1.7. DEGs enriching to this GO term and upregulated following infection by the ancestral and B.1.1.7 variants only were part of T cell activation pathways (e.g., *IFNG, IL12RB1, TBX21, XCL1*) (Fig. S2B).

Other DEGs that were upregulated following infection with all three variants enriched to GO term “blood vessel development”. These genes are involved in angiogenesis (e.g., *ANGPTL2, ANGPTL4, ADM2, HOX1*), apoptosis (e.g., *BAK1, FASLG*), tissue remodeling (e.g., *CHI3L1, MMP19*), and leukocyte chemotaxis (e.g., *CCL11, CCL2, CXCL10, CXCL17*) (Fig. S2C). Regulators of angiogenesis, like *SOX4* and *KDR*, and genes involved in tissue remodeling (e.g., ADAM12, SHH) were downregulated only in infections with the B.1.1.7 and ancestral virus (Fig. S2C). Shared DEGs that enriched to GO term “lymphocyte activation” included genes important for B cell maturation (e.g., *AIRE, CD27, CD38, ICOS, TNFSF13B*) as well as negative regulation of T cell responses (e.g., *CD274, CTLA4, FOXP3, IDO1, PDCDC1, PDCD1LG2*) (Fig. S2E). DEGs shared between B.1.1.7- and ancestral-infected hamsters were important for T cell activation (e.g., *PRKCQ, TNFSF9*), cytotoxic responses (e.g., *KLRK1, PRF1*) myeloid cell activation (e.g., *IFNG, SLAMF1, CD177, CXCL6*) and IL-6 production (e.g., *TLR1, IL-6, IL18RAP, C3AR1, C1QA*) (Fig. 4E, S2D).

We next analyzed DEGs unique to each infection to understand infection-specific transcriptional responses (Fig. 5). The largest group of unique DEGs was detected following B.1.1.7 infection (n=648). These unique DEGs enriched to GO terms reflecting tissue remodeling (e.g., “response to growth factor”, “tissue morphogenesis”) (Fig. 5A). Most DEGs in these GO terms are downregulated and associated with angiogenesis (e.g., *ENG, JCAD, PDFGB, VEGFD*) and lung development (e.g., *FZD1, SOX17, TMEM100, VANGL2*), while a smaller upregulated portion was associated with cell death (e.g., *APAF1, CASP3*), and protein degradation (e.g., *CASP3, DAB2, SFRP1*). Other DEG enriched to GO terms associated with host defense (e.g., “adaptive immune response”) and cell recruitment (e.g., “chemotaxis”) were identified. Most of these DEGs were upregulated and are important for antigen presentation (e.g., *CD74, HLA-DRA, B2M*) and natural killer (NK) cell-mediated immunity (e.g., *CD84, IL12A*) (Fig. 5B-D). Notable DEGs unique to infection with B.1.351 play a role in cell morphogenesis (e.g., *ACTA2, ACTC1, FGF1*), myeloid cell differentiation (e.g., *CAV3, PDE1B, TFRC*), and response to injury (e.g. *COL4A3, MPL, TSPAN*) (Fig. 5E). Downregulated DEGs unique to infection with the ancestral strain encoded components of cellular respiration (e.g., *MT-C03, MT-ND1*) and mediators of cell adhesion (e.g., *IKF26B, VIT*) (Fig. 5F).

**Fig. 5.**
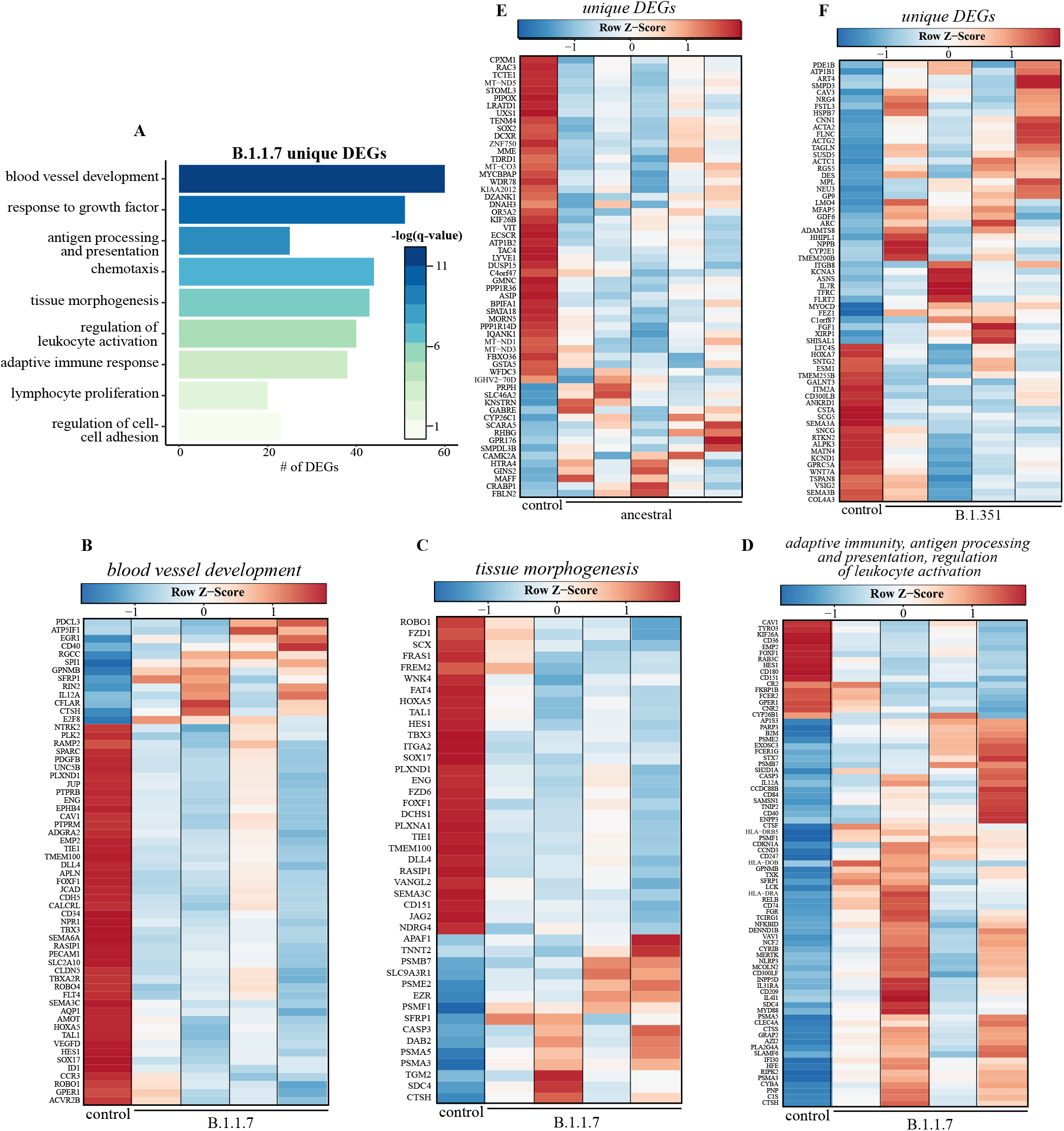
Transcriptional response unique to B.1.1.7 variant suggests distinct host responses. **(A)** Functional enrichment of DEGs unique to B.1.1.7 variant infection at 4 DPC (n=684). Horizontal bars represent the number of genes enriching to each GO term with color intensity representing the negative log of the FDR-adjusted p-value [-log(q-value)]. Range of colors based on the GO terms with the lowest and highest –log(q-value) values. Heatmaps representing B.1.1.7 variant unique DEGs enriching to GO terms from panel A: **(B)** “blood vessel development”, **(C)** “tissue morphogenesis” and **(D)** adaptive terms “adaptive immunity”, “antigen processing and presentation”, and “regulation of leukocyte activation.” Heatmaps of DEGs unique to **(E)** Ancestral (n=58) and **(F)** B.1.351 variants (n=64) at 4 DPC. Columns of all heatmaps represent the average rpkm of controls and rpkm of a single variant-infected animal. Range of colors per each heatmap is based on scaled and centered rpkm values of the represented DEGs. Red represents upregulation; blue, downregulation.

### Digital cell quantification in hamster lungs

Since Syrian golden hamsters lack adequate reagents for immunophenotyping, we performed digital cell quantification (DCQ) to predict changes in immune cell populations using the IRIS immune cell database^18^. Changes in gene expression were predicted to be associated with increased frequencies of activated NK cells, activated dendritic cells (DCs), and neutrophils after ancestral and B.1.1.7 infection (Fig. S3A). In contrast, B.1.351 infection was associated with a decrease in NK cells and monocytes (Fig. S3B). Reduced frequencies of B cells were predicted for all infections, while increases in Th1 and Th2 CD4+ T cells were only predicted after B.1.1.7 infection (Fig. S3B).

## Discussion

Over the last several months a number of SARS-CoV-2 VOC have emerged. These VOC are associated with increased transmissibility and enhanced viral fitness due to mutations in S. Several studies have shown that the N501Y mutation harborded in both the B.1.1.7 and the B.1.351 variants utilized here increases ACE2 binding and enhances transmission capabilities^4,5,19^. The K417N and E484K mutations introduced into the S of the B.1.351 enhances the ability to evade pre-existing humoral responses^3,7,10,20–22^. A comparative study of viral pathogenesis of VOC has recently been conducted in the hamster model^23^. The study measured the viral burden, histopathology, and select cytokine gene expression induced by VOC compared to the prototypic Wuhan-Hu-1 isolate and an isolate harboring the secondary D614G mutation in S. The study showed no significant differences in viral burden and histopathologic findings in the hamster lungs at 4 DPC, but enhanced expression of cytokine genes was described in hamsters infected with the B.1.1.7 variant.^23^ However, longitudinal analysis of the host response to VOC and the degree of cross-protection is lacking. Therefore, in this study, we sought to evaluate the longitudinal impact of these VOC on the host immune and transcriptional responses.

Syrian golden hamsters were chosen for this study as they are highly susceptible to infection and were found to have high viral replication in the lungs. Hamsters were infected IN with the ancestral, B.1.1.7 or B.1.351 variants. Challenged hamsters displayed moderate weight loss lethargy, rapid breathing, and ruffled fur, but were able to clinically recover by 14 DPC as previously described^14,24,25^. As recently reported, no discernable differences in gross pathology or lung viral burden were noted among all three groups^26^. However, B.1.1.7 sgRNA persisted longer in the hamster lungs. Analysis of the viral genomes recovered post-infection showed no changes in the ancestral- and B.1.1.7-infected hamsters; however, we detected three mutations in all B.1.351-infected animals. The two nonsynonymous mutations occurred in nsp3 and ORF3a, both of which have been implicated in evasion of type I IFN^27,28^. A second mutation in nsp3 was also identified in a single B.1.351-infected animal. The implication of these mutations remains to be elucidated.

Analysis of the humoral response revealed that the overall IgG response of the infected hamsters did not result in robust differences amongst the variants; however, the neutralization cross-protection depended on the variant the hamster was initially exposed to. Specifically, infection with B.1.1.7 results in the widest breath of neutralization activity despite comparable binding antibody titers. This phenomenon was most noticeable at 14 DPC, and was still evident 28 DPC when the humoral response is more mature. Moreover, the overall neutralization activity, regardless of initial challenge virus, against B.1.351 is much lower than the other two variants, suggesting that B.1.351 may have indeed an enhanced ability to evade humoral immune responses. The overall IgG response of the infected hamsters did not result in robust differences amongst the variants; however, the neutralization cross-protection depended on the variant the hamster was initially exposed to. Our data demonstrates that early in the humoral response (14 DPC) antibodies induced by B.1.1.7 infection show an increased crossrreacitvity compared to the other variants tested. By 28 DPC, when the humoral response is more mature, this differences is less prominent, but the trend remains the same. This observation suggests that the timing of the antibody response could affect the crossreactivity potential. Notably, the neutralization capacity of crossreactive antibodies and homologous antibodies against B.1.351 is much lower than that of the other two variants tested. This observation is reflective of previous studies that attribute increased antibody evasion to this VOC^3,4,8–10,12,21,22^, demonstrating that the hamster model reflects the differences in humoral responses and effectivity of prior immunity seen in clinical cases^29^.

A significant challenge when using the hamster model is the lack of reagents to analyze cellular immune responses^30–36^. Therefore, we employed transcriptomic analysis to elucidate differences in the host responses to VOC compared to the ancestral variant in the lungs of hamster 4 DPC, as has been done for other studies ^37–39^. Our transcriptional analysis of lung tissues at peak infection identified distinct, but also overlapping transcriptional signatures for each variant. All infections exhibited gene expression patterns associated with innate antiviral responses, notably type I IFN signaling, mobilization of lymphocytes, and apoptosis^40–43^. The type I IFN response is critical for rapid control of viral infection^44^. However, dysregulated innate immune and type I IFN responses can result in tissue damage and oxidative stress as noted in other viral infections, including influenza virus and Ebola virus in addition to severe COVID-19^40,41,45,46^. Our data differs from those reported in studies where a suppressed IFN response in the peripheral blood, the bronchoalveolar lavage, and lungs obtained at autopsy from individuals with severe COVID-19^47–52^. A potential explanation for this difference is the fact that we profiled the lungs during the peak of viral replication and virus-induced pathology (4 DPC) while clincal cases rarely present viral antigen at the time of death, rather immune dysregulation and coagulation abnormalities are the casue of death^53–55^. Additionally, the Syrian golden hamster model does not mimic severe COVID-19 intersitial pneumonia in that clinical symptomology is less severe and none of the animals in this model succumb to disease. Interestingly, transcriptional inflammatory indicators were particularly heightened following infection with B.1.1.7 and least severe following infection with B.1.351. Expression of several inflammatory and complement genes were only upregulated following infection with B.1.1.7 and ancestral variants, while *NFκB1* was upregulated only following infection with B.1.1.7^56,57^. *In vitro* and *in vivo* NFκB-driven inflammatory responses have been previously associated with severe COVID-19^48,50,58,59^. Additionally, NK cell activation was evident by higher expression of cytolytic molecules (e.g., *PRF1*). This inflammatory damage facilitates immune cell influx, including inflammatory cells like neutrophils, which we predicted to increase in all infections^48^. Moreover, significant increases in IL-2-stimulated NK cells was also predicted following infection with the ancestral and B.1.1.7 variants. Expression of canonical T cell regulatory and exhaustion markers like *CTLA4, CD274* (PD-L1), and *FOXP3* suggests compensatory mechanisms to reduce tissue damage.

Transcriptional changes were also predicted to result in significant B cell loss in the lungs following infection with all three viruses. Previous studies indicate that B cell lymphopenia does not preclude robust antibody responses^60–62^. This re-distribution could indicate B cell migration to lymphoid tissue for priming. Indeed, significant neutralizing and binding antibody titers were detected following all three infection, albeit lower following infection with B.1.135.

Furthermore, we detected a large number of DEGs related to tissue morphogenesis and angiogenesis in all infections^63,64^. Microvascular injury can further exacerbate inflammation-driven lung fibrosis^65^. Additionally, genes that play a role in tissue repair were downregulated following infection with the B.1.351 and ancestral variants.

In this study we describe the pathogenesis of the SARS-CoV-2 variants and the development of crossreactive neutralizing antibodies. To our knowledge this is the first study performing a comparative and longitudinal analysis of the antibody response after SARS-CoV-2 VOC infection. Our data show that infection with the B.1.1.7 VOC results in a broader antibody response compared to infection with B.1.35 VOC. This broader response could be in part mediated by the more robust transcriptional response elicited by this variant that includes a larger induction of antiviral and inflammatory pathways. Future experiments should assess transcriptional changes beyond 4 DPC to determine the kinetics of the host response at this critical site. Moreover, additional studies should investigate the mechanisms by which the mutations detected in the B.1.35 VOC lead to reduced neutralization potential.

## Methods

### Ethics statement

All infectious work with SARS-CoV-2 was performed in the containment laboratories at the Rocky Mountain Laboratories (RML), Division of Intramural Research, National Institute of Allergy and Infectious Diseases, National Institutes of Health. RML is an institution accredited by the Association for Assessment and Accreditation of Laboratory Animal Care International (AAALAC). All procedures followed standard operating procedures (SOPs) approved by the RML Institutional Biosafety Committee (IBC)^66^. Animal work was performed in strict accordance with the recommendations described in the Guide for the Care and Use of Laboratory Animals of the National Institute of Health, the Office of Animal Welfare and the Animal Welfare Act, United States Department of Agriculture. The studies were approved by the RML Animal Care and Use Committee (ACUC). Procedures were conducted in animals anesthetized by trained personnel under the supervision of veterinary staff. All efforts were made to ameliorate animal welfare and minimize animal suffering; food and water were available *ad libitum*.

### Cells and Viruses

VeroE6 cells were grown at 37°C and 5% CO_2_ in Dulbecco’s modified Eagle’s medium (DMEM) (Sigma-Aldrich, St. Louis, MO) containing 10% fetal bovine serum (FBS) (Wisent Inc., St. Bruno, Canada), 2 mM L-glutamine (Thermo Fisher Scientific, Waltham, MA), 50 U/mL penicillin (Thermo Fisher Scientific), and 50 μg/mL streptomycin (Thermo Fisher Scientific). SARS-CoV-2 ancestral isolate nCoV-WA1-2020 (MN985325.1)^67^, SARS-CoV-2 isolate B.1.351 (hCoV-19/South African/KRISP-K005325/2020), or SARS-CoV-2 isolate B.1.1.7 (hCOV_19/England/204820464/2020) were used for the neutralizing antibody assays. The following reagent was obtained through BEI Resources, NIAID, NIH: Severe Acute Respiratory Syndrome-Related Coronavirus 2, Isolate hCoV-19/England/204820464/20200, NR-54000, contributed by Bassam Hallis. SARS-CoV-2 B. 1.351 was obtained with contributions from Dr. Tulio de Oliveira and Dr. Alex Sigal (Nelson R Mandela School of Medicine, UKZN). All viruses were grown and titered on Vero E6 cells, and sequence confirmed.

### Animal study

Fifty female Syrian golden hamsters (5-8 weeks of age) were used in this study^14^. Five animals were used as uninfected controls; three study cohorts for challenge with the ancestral virus and variants B1.1.7 and B.1.351 consisted of 15 hamsters each. On day 0, hamsters were infected with SARS-CoV-2 as previously described ^14^. On 4, 14 and 28 DPC, 5 hamsters per group were euthanized for sample collection.

### RNA extraction and RT-qPCR

RNA from blood and oral swab samples was extracted using the QIAamp Viral RNA Mini Kit (QIAGEN) according to manufacturer specifications. Lung tissue, a maximum of 30 mg each, was processed and RNA was extracted using the RNeasy Mini Kit (QIAGEN) according to manufacturer specifications. One step RT-qPCR for genomic viral RNA was performed using specific primer-probe sets and the QuantiFast Probe RT-PCR +ROX Vial Kit (QIAGEN), in the Rotor-Gene Q (QIAGEN) as described previously ^68^. Five μL of each RNA extract were run alongside dilutions of SARS-CoV-2 standards with a known concentration of RNA copies.

### Enzyme-linked immunosorbent assay

Serum samples from SARS-CoV-2-infected animals were inactivated by gamma-irradiation and used in BSL2 according to IBC-approved SOPs. NUNC Maxisorp Immuno plates were coated with 50 μl of 1 μg/mL of recombinant SARS-CoV-2 S (S1+S2) antigen at 4°C overnight and then washed three times with PBS containing 0.05% Tween 20 (PBST). The plates were blocked with 3% skim milk in PBS for 1 hour at room temperature, followed by three additional washes with PBST. The plates were incubated with 50 μl of serial dilutions of the samples in PBS containing 1% skim milk for 1 hour at room temperature. After three washes with PBST, the bound antibodies were labeled using 50 μl of 1:2,500 peroxidase anti-hamster IgG (H+L) (SeraCare Life Sciences) diluted in 1% skim milk in PBST. After incubation for 1 hour at room temperature and three washes with PBST, 50 μl of KPL ABTS peroxidase substrate solution mix (SeraCare Life Sciences) was added to each well, and the mixture was incubated for 30 min at room temperature. The optical density (OD) at 405 nm was measured using a GloMax® explorer (Promega). The OD values were normalized to the baseline samples obtained with naïve hamster serum and the cutoff value was set as the mean OD plus standard deviation of the blank.

### Virus neutralization assay

The day before this assay, VeroE6 cells were seeded in 96-well plates. Serum samples were heat-inactivated for 30 min at 56°C, and 2-fold serial dilutions were prepared in DMEM with 2% FBS. Next, 100 TCID_50_ of SARS-CoV-2 were added and the mixture was incubated for 1 hour at 37°C and 5% CO_2_. Finally, media was removed from cells and the mixture was added to VeroE6 cells and incubated at 37°C and 5% CO_2_ for 6 days. Then the cytopathic effect (CPE) was documented, and the virus neutralization titer was expressed as the reciprocal value of the highest dilution of the serum which inhibited virus replication (no CPE).

### Histology and immunohistochemistry

Necropsies and tissue sampling were performed according to IBC-approved SOPs. Tissues were fixed in 10% neutral buffered formalin with two changes, for a minimum of 7 days. Tissues were placed in cassettes and processed with a Sakura VIP-6 Tissue Tek, on a 12-hour automated schedule, using a graded series of ethanol, xylene, and ParaPlast Extra. Embedded tissues are sectioned at 5 μm and dried overnight at 42°C prior to staining. Specific anti-CoV immunoreactivity was detected using Sino Biological Inc. SARS-CoV/SARS-CoV-2 nucleocapsid antibody (Sino Biological cat#40143-MM05) at a 1:1000 dilution. The secondary antibody was the Vector Laboratories ImPress VR anti-mouse IgG polymer (cat# MP-7422). The tissues were then processed for immunohistochemistry using the Discovery Ultra automated stainer (Ventana Medical Systems) with a ChromoMap DAB kit (Roche Tissue Diagnostics cat#760–159). All tissue slides were evaluated by a board-certified veterinary pathologist and a pathology scored was assigned based on the following observations; 0= no pathology, 1= minimal, 2= mild, 3=moderate, 4= severe (Fig. S1C).

### cDNA library construction and sequencing

Quality and quantity of RNA lung samples at 4 DPC were determined using an Agilent 2100 Bioanalyzer. cDNA libraries were constructed using the NEB Next Ultra II Direction RNA Library Prep Kit (Thermo Fischer). RNA was treated with RNase H and DNase I following depletion of ribosomal RNA (rRNA). Adapters were ligated to cDNA products and the subsequent ^~^300 base pair (bp) amplicons were PCR-amplified and selected by size exclusion. cDNA libraries were assessed for quality and quantity prior to 150 bp single-end sequencing using the Illumina NovaSeq platform.

### RNA-Seq Bioinformatic analysis

Preliminary data analysis was performed with RNA-Seq workflow module of systemPipeR, developed by Backman *et. al*^69^. RNA-Seq reads were demultiplexed, quality-filtered and trimmed using Trim Galore (average Phred score cut-off of 30, minimum length of 50 bp).

FastQC was used to generate quality reports. Hisat2 was used to align reads to the reference genome *Mesocricetus auratus* (Mesocricetus_auratus.MesAur1.0.dna.toplevel.fa) and the Mesocricetus_auratus.MesAur1.0.103.gtf file was used for annotation. Raw expression values (gene-level read counts) were generated using the summarizeOverlaps function and normalized (read per kilobase of transcript per million mapped reads, rpkm) using the edgeR package. Statistical analysis with edgeR was used to determine differentially expressed genes (DEGs) meeting the following criteria: genes with median rpkm of ≥1, a false discovery rate (FDR) corrected p-value ≤ 0.05 and a log_2_fold change ≥1 compared to control tissues.

Functional enrichment of DEGs was performed using Metascape to identify relevant GO terms^70^. Digital cell quantification was performed using ImmQuant with the IRIS database. Heatmaps, bubbleplots, Venn diagrams and violin plots were generated using R packages ggplot2 and VennDiagrams. Graphs were generated using GraphPad Prism software (version 8).

### SARS-CoV-2 viral genome library construction and sequencing

Enrichment of SARS-CoV-2 was performed using the Qiagen QIASeq SARS-CoV-2 Primer Panel (V.2). Libraries were constructed from resulting SARS-CoV-2 amplicons using the Qiagen QIASeq FX DNA Library preparation kit. Briefly, adapters were ligated to cDNA products and the ^~^300 bp amplicons were minimally PCR-amplified. cDNA libraries were assessed for quality and quantity prior to 150 bp paired-end sequencing using the Illumina HiSeq platform (≥ 1 M reads per sample).

### SARS-CoV-2 viral genome assembly and bioinformatic analysis

Reads were demultiplexed and quality-filtered using Trim Galore (average Phred score cut-off of 30, minimum length 100 bp). FastQC was used to generate quality reports. MaskPrimers.py from the pRESTO R package was used to remove primers prior to alignment to the SARS-CoV-2 genome using BWA-mem software version 0.7.17. The following reference genomes were used for Ancestral, B.1.1.7 and B.1.351 variants: WA_MN985325.1, EPI_ISL_683466, and EPI_ISL_6786156. All genomes had greater than 95% coverage and 10X depth. Single nucleotide polymorphisms and amino acid changes were identified using CorGAT.

### Statistical analyses

All statistical analysis was performed in Prism 8 (GraphPad). Two-tailed Mann-Whitney test was conducted to compare differences between groups for data in Figs. 2, 3 and S1. Statistical significance was determined using one-way ANOVA with multiple comparisons for the bioinformatic analysis with comparisons made among variant- and control-challenged animals Statistically significant differences are indicated as follows: p<0.0001 (****), p<0.001 (***), p<0.01 (**) and p<0.05 (*).

## Data Availability

All transcriptomic sequencing data are accessible at BioProject PRJNAXXXX upon publication.

## Acknowledgments

We thank members of the Molecular Pathogenesis Unit, Virus Ecology Section, and Research Technology Branch (all NIAID) for their efforts to obtain and characterize the SARS-CoV-2 isolates. We also thank the Rocky Mountain Veterinary Branch, NIAID for supporting the animal studies, and Anita Mora (NIAID) for assistance generating the pathology figures.

## Author contributions

A.M. conceived the idea and secured funding. K.L.O., I.M, and A.M. designed the studies. K.L.O., C.S.C., K.S., T.G., P.F., and A.M. conducted the animal studies, processed the samples and acquired the data. A.N.P. performed transcriptomics analysis and viral genome sequencing. K.L.O., A.N.P., C.S.C., I.M., and A.M. analyzed and interpreted the data. K.L.O., A.N.P., I.M., and A.M. prepared the manuscript with input from all authors. All authors approved the manuscript.

---

**Figure S1.**
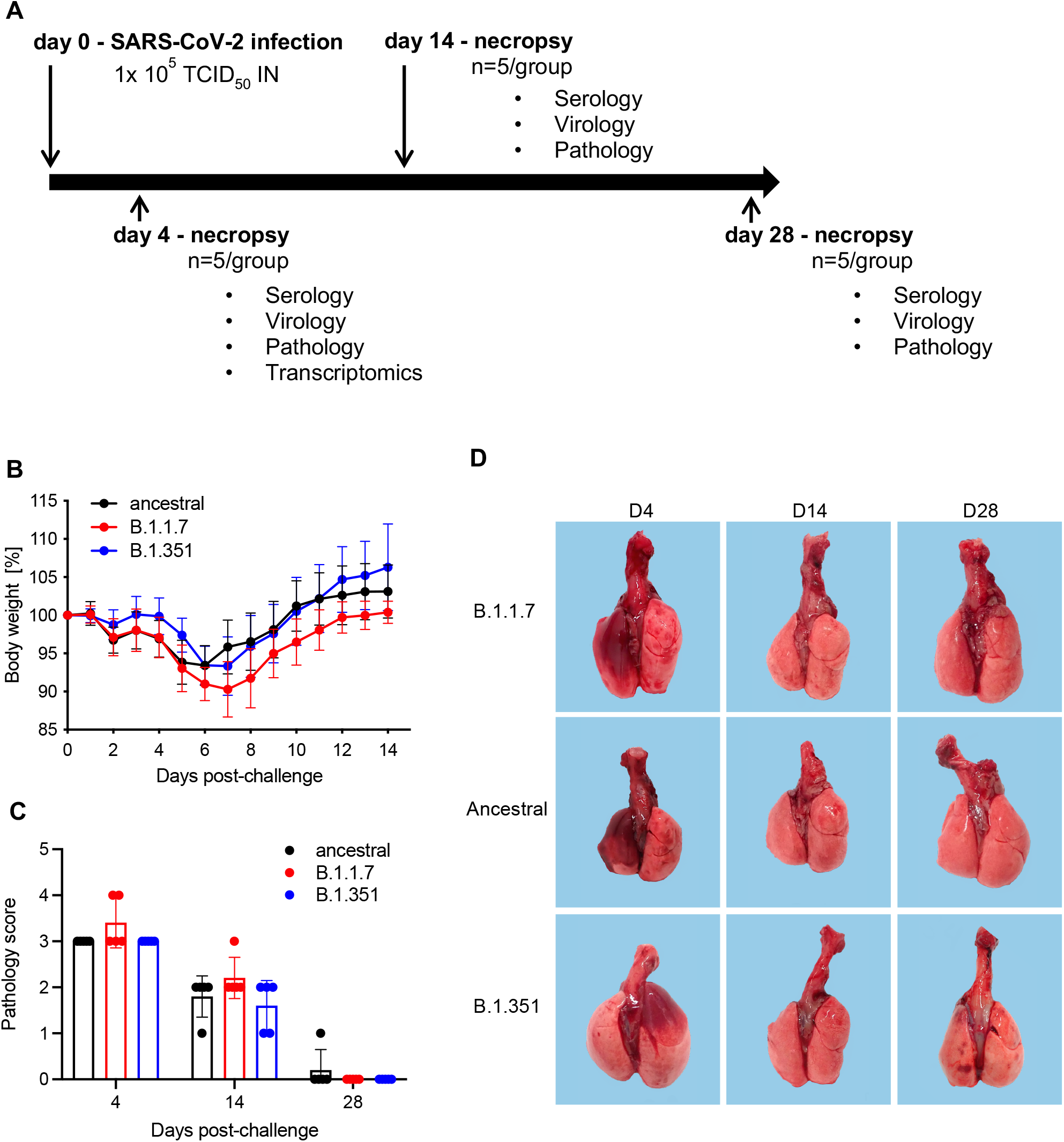
Study outline, body weight changes and pathology of infected hamsters. **(A) Schematic study outline.** (B) Body weight changes in hamsters (n=10/group). (C) Evidence of interstitial pneumonia was recorded in histopathology samples. (D) Representative pictures of hamster lungs with lesions during disease progression. Gross lung images at day (D) 4, 14 and 28 post-challenge.

**Figure S2.**
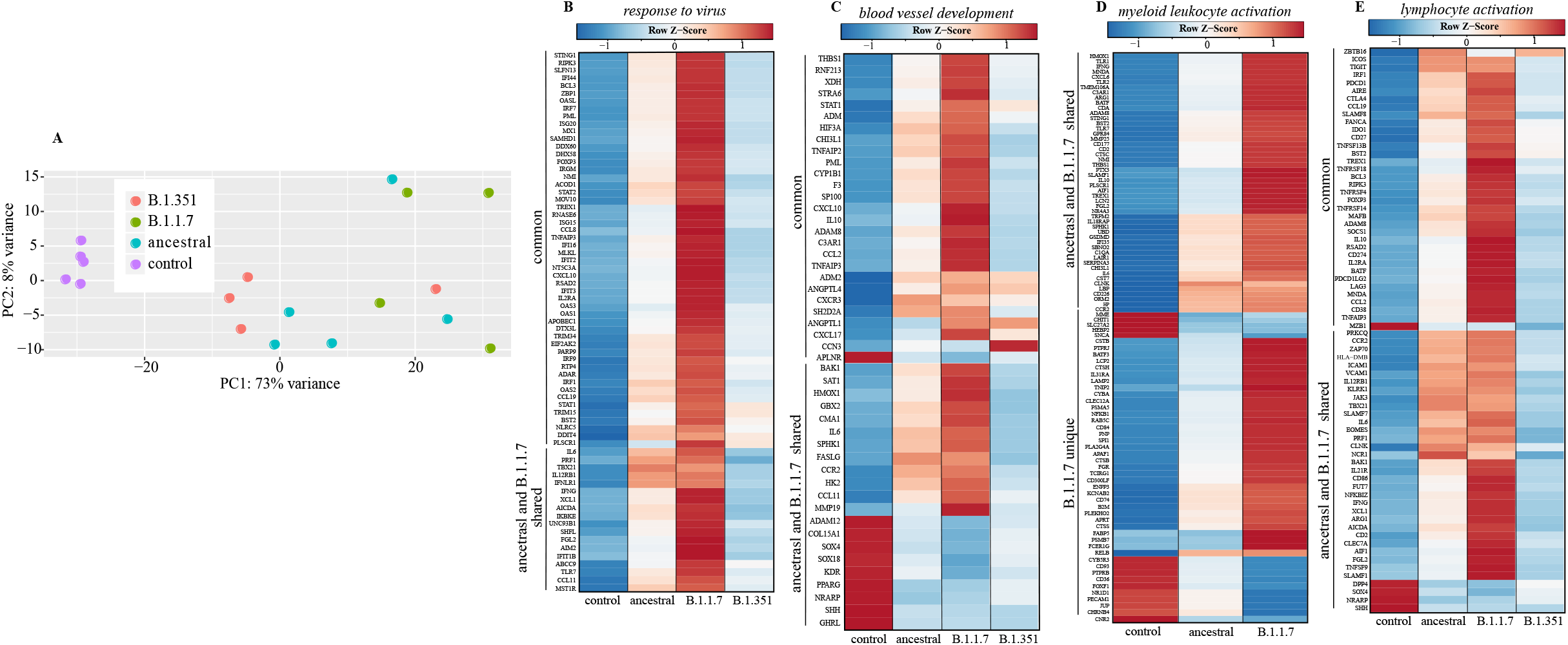
SARS-CoV-2 variants induce distinct transcriptional changes. (A) Principal component analysis of control (n=5) animals and infected animals 4 DPC with ancestral (n=5), B.1.1.7 (n=4) or B.1.351 (n=4) variants. Heatmaps representing DEGs enriching to GO terms from Fig. 5E including (B) “response to virus”, (C) “blood vessel development”, (D) “myeloid leukocyte activation” and (E) lymphocyte activation.” DEGs are either shared among all variant infections or between ancestral and B.1.1.7 variant infections. Columns of all heatmaps represent the average rpkm. Range of colors per each heatmap is based on scaled and centered rpkm values of the represented DEGs. Red represents upregulation; blue, downregulation.

**Figure S3.**
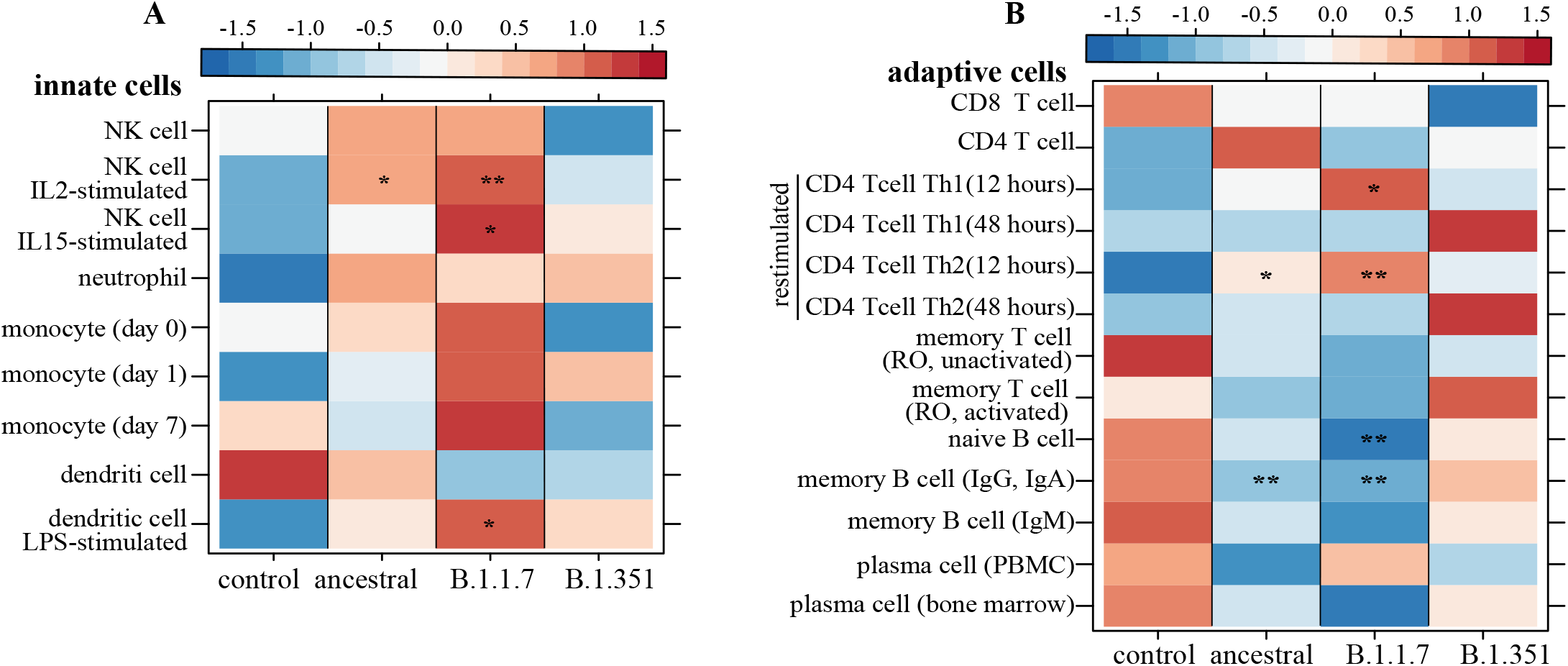
Digital cell quantification in hamster lungs. Heatmaps representing relative changes in (A) innate and (B) adaptive immune cell frequencies using ImmQuant with IRIS database. Each column represents the average relative expression level of the given immune cell. Range of colors per each heatmap is based on scaled and centered relative expression values. Red represents upregulation; blue represents downregulation. Statistical significance is indicated **p < 0.01 and *p < 0.05.

